# Heterogeneous Impacts of Protein-Stabilizing Osmolytes on Hydrophobic Interaction

**DOI:** 10.1101/328922

**Authors:** Mrinmoy Mukherjee, Jagannath Mondal

## Abstract

Osmolytes’ mechanism of protecting proteins against denaturation is a longstanding puzzle, further complicated by the complex diversities inherent in protein sequences. An emergent approach in understanding osmolytes’ mechanism of action towards biopolymer has been to investigate osmolytes’ interplay with hydrophobic interaction, the major driving force of protein folding. However, the crucial question is whether all these protein-stabilizing osmolytes display a *single unified* mechanism towards hydrophobic interactions. By simulating the hydrophobic collapse of a macromolecule in aqueous solutions of two such osmoprotectants, Glycine and Trimethyl N-oxide (TMAO), both of which are known to stabilize protein’s folded conformation, we here demonstrate that these two osmolytes can impart mutually contrasting effects towards hydrophobic interaction. While TMAO preserves its protectant nature across diverse range of polymer-osmolyte interactions, glycine is found to display an interesting cross-over from being a protectant at weaker polymer-osmolyte interaction to a denaturant of hydrophobicity at stronger polymer-osmolyte interactions. A preferential-interaction analysis reveals that a subtle balance of *conformation-dependent* exclusion/binding of osmolyte molecules from/to the macromolecule holds the key to overall heterogenous behavior. Specifically, TMAO’s consistent stabilization of collapsed configuration of macromolecule is found to be a result of TMAO’s preferential binding to polymer via hydrophobic methyl groups. However, polar Glycine’s cross-over from being a protectant to denaturant across polymer-osmolyte interaction is rooted in its switch from preferential exclusion to preferential binding to the polymer with increasing interaction. Overall, by highlighting the complex interplay of osmolytes with hydrophobic interaction, this work puts forward the necessity of quantitative categorization of osmolytes’ action in protein.

## Introduction

Osmoprotectants, which are small cosolutes, stabilize the folded conformation of proteins by counteracting the action of a denaturant, such as urea and help organisms cope up with the osmotic stress. ^1–5^ One of the most well-studied protecting osmolytes in this context, Trimethyl N-oxide (TMAO), has been at the center of multiple investigations due to its stabilising role on protein folding as an osmolyte.^6–13^ Apart from TMAO, two other osmoprotectants, namely, glycine and glycine betaine have also emerged as key chemical chaperones of interest for their stabilizing role of protein’s folded conformation against denaturation.^14–19^ In case of proteins, it has now been established that all these osmoprotectants counteract the denaturing effect of urea towards protein. One of the most prevalent hypotheses in the counteraction role of both TMAO^20–24^ and glycine or glycine betaine^14–18^ has been the preferential exclusion of cosolutes from the surface of the protein. The objective of the current work is to show that the uniform principle of stabilization by so-called preferential exclusion of osmolytes does not quite hold together across all osmoprotectants in case of hydrophobic interaction, the major driving force underlying protein stability.

The action of these osmolytes on the hydrophobic interaction has only started to garner significant attention recently, with TMAO being the key osmolyte at the forefront of interest. In a clear departure from the prevalent perception of preferential exclusion mechanism by the protecting osmolytes, Mondal et al^25^ earlier proposed the idea of stabilization of collapsed conformation of a hydrophobic macromolecule by aqueous solution of TMAO via preferential binding to the polymer surface. By computationally simulating a model hydrophobic polymer and by utilizing Wyman-Tanford preferential binding theory,^26,27^ Mondal et al showed that, while TMAO preferentially binds to the polymer surface, it is the relative difference in the extent of preferential binding of TMAO on collapsed conformation compared to that on an extended conformation which guides the direction of conformational equilibrium of a hydrophobic polymer. This hypothesis of preferential binding of TMAO on a model hydrophobic polymer system was subsequently experimentally validated on a synthetic hy-drophobic polymer namely polystyrene.^28^ Since then, the idea of stabilization of collapsed conformation of macromolecule via preferential binding of TMAO on the surface, in contrast to preferential exclusion, is slowly getting recognized in multiple new studies on hydrophobic polymer^29–35^ and hydrophobic peptide^19^ as well. Recently, by combining experiments and MD simulations of a hydrophobic peptide in aqueous solution of three different osmolytes, namely TMAO, glycine and glycine betaine, Noid and Cremer group^19^ showed that while the globular conformation of hydrophobic peptide can get stabilized by these osmolytes, the mechanism of action of TMAO on this peptide was found to be quite different from that of glycine and betaine: TMAO was found to accumulate near the peptide surface, while glycine and betaine were found to preferentially exclude from the peptide surface. A similar observation of preferential accumulation of TMAO on protein surface was also made by Dias and coworkers^36^ in simulations of poly-leucine, Trp-cage protein and amyloids. These recent observations of preferential accumulation of TMAO on a peptide surface are quite distinct from past osmolyte-based studies on proteins and have been majorly consistent with the past observations of TMAO’s preferential binding on hydrophobic surface.^25,28^

The aforementioned discussion raises the question whether all protecting osmolytes’ action on a hydrophobic (bio)macromolecule is characterized by a *single unified* mechanism, similar to that of TMAO. An answer to this question can potentially lead to perception of an unified underlying mechanism of action by all osmoprotectants towards hydrophobic effect. The current article, by quantitatively assessing the role of the protein-stabilizing osmolytes, namely glycine (figure 1A) and TMAO (figure 1B), towards collapse behavior of hydrophobic polymer, would show that the overall pictures are quite heterogenous: Depending on polymer-osmolyte interactions, these osmolytes can even exert mutually similar or opposite effect on hydrophobic interaction. We address this question by independently simulating the effect of TMAO and glycine on collapse behavior of a 32-bead uncharged hydrophobic polymer and subsequently interpreting the results using Wyman-Tanford preferential binding theory.^26,27^ This polymer chain has been previously studied by Mondal et al^25,37^ extensively and the details of the models can be found else-where.^25,37^ The associated osmolyte and water forcefields are detailed in SI text. All simulations are individually performed using 0.5 M aqueous solution of glycine and 0.5 M aqueous solution of TMAO (we have also performed simulations with aqueous solution of glycine betaine, the result of which will be briefly summarized in SI). A representative snapshot of starting configuration is presented in figure 1 C and D. As would be seen later, the polymer of our interest being devoid of any complexity, allows one to simulate the effect of diverse range of polymer-osmolyte interactions towards hydrophobic interaction.

**Figure 1:**
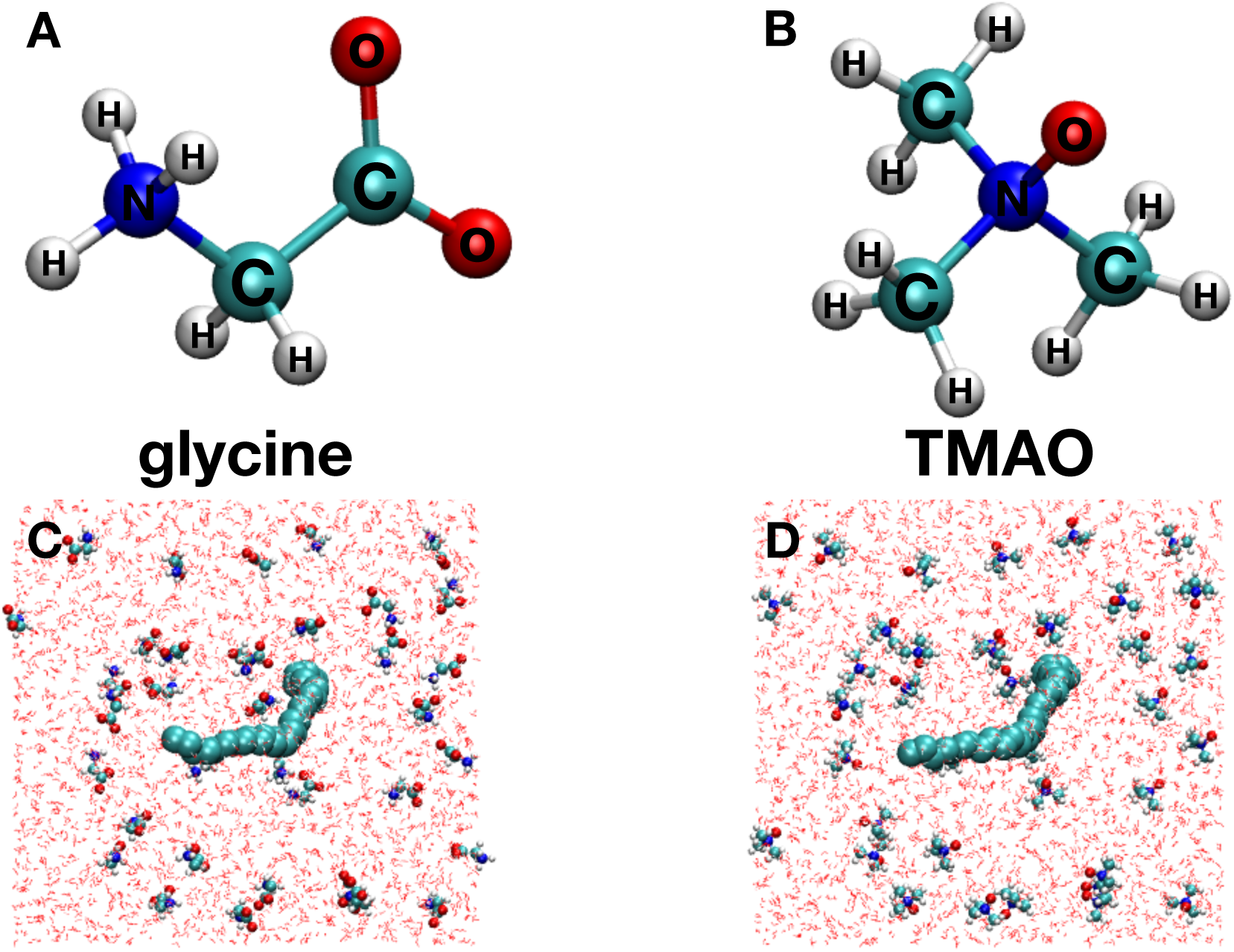
A representative (A) glycine and (B) TMAO molecule and a representative configuration of extended polymer in 0.5 M aqueous mixture of (C) glycine and (D) TMAO (water molecules are represented by red lines).

## Method and Model

*Simulation model:* We have simulated a system of a 32-bead polymer chain in (see Figure 1 in main text) two different aqueous media of osmolytes, namely glycine and TMAO. The polymer chain has been previously studied by Mondal et al^25,37^ extensively and the details of the models can be found else-where.^28,37^ Here we provide a brief overview of the models. We have used SPC/E^38^ model for water, CHARMM27^39^ force field for glycine and for TMAO the force fields developed by Shea and co workers.^24,40^ In our previous works,^25,32^ we have explored multiple forcefields of TMAO and based on the results, we have decided to zero in on that developed by Shea and coworkers.^24,40^ In this work, all weak dispersion nonbonding interactions have been modeled by Lennard Jones (LJ) interactions and geometric combination rules (detailed later) have been applied for all intermediate interactions. Since the current work aims to dissect the role of the osmolytes on hydrophobic polymer collapse, we mainly explore the effect of variation of polymer-cosolute interaction on stabilization of polymer-collapse at fixed intra-polymer and polymer-water dispersion interaction. The polymer’s internal bead-bead dispersion interaction (*∊*_*b*_) has been kept fixed at 1.0 kJ/mol and polymer-water dispersion interaction is kept constant by fixing *∊*_*p*_ to 1.0 kJ/mol via combination rule 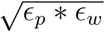 where *∊*_*w*_ is the LJ interaction of oxygen atoms of SPC/E water model. As we will see later, the intra-bead dispersion interaction of 1 kJ/mol within the polymer is sufficient to induce a hydrophobic polymer collapse. The polymer-osmolyte interaction is defined by 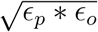 where *∊*_*o*_ represents the interaction of different osmolytes atoms. For each osmolyte solution, 3 different cases (hence a total of six cases for glycine and TMAO) were simulated by varying the *∊*_*p*_ in polymer-osmolyte interactions from 0.6 to 1.4 kJ/mol at an interval of 0.4 kJ/mol.

All simulations are individually performed at 0.5 M aqueous solution of glycine and 0.5 M aqueous solution of TMAO. This osmolyte concentration is very close to what was earlier employed by Cremer and co-workers^19^ (0.55 M) in their theory/experimental investigations of these two osmolytes on a protein. In our simulations, these are achieved by incorporating the polymer in a solution of 37 osmolyte (glycine or TMAO, depending on the case) molecules and 3888 water molecules in an cubic box of initial dimension 5 *x* 5 *x* 5 nm. The concentrations of osmolytes are individually maintained at ≈ 0.5M by adjusting the box at an average temperature of 300 K and average pressure of 1 bar using NPT ensemble.

As an aside, we have simulated the effect of aqueous solution of glycine betaine on the hydrophobic polymer using the similar approach as detailed for TMAO and glycine. For the sake of brevity, we have summarized the result involving glycine betaine in the supporting information.

*Method:* The work assesses the relative stability of collapsed conformation of the polymer compared to extended or unfolded conformation by quantifying the underlying conformational free energy landscape of the polymer along the radius of gyration of the polymer in each of the aforementioned osmolyte solutions. Towards this end, we have performed umbrella sampling simulations^41^ using radius of gyration of the polymer as a collective variable. To generate the representative configurations required for each of the so-called ‘umbrella sampling windows’, we first performed independent equilibrium Molecular Dynamics simulations for 100 ns at constant pressure of 1 bar and temperature of 300 K, starting with an extended all-trans configuration of the polymer. To avoid any bias and to supplement any missing configurations for subsequent umbrella sampling simulations, these simulations were also repeated starting with a collapsed configuration of the polymer.

For umbrella sampling simulations, the values of the radius of gyration ranged from 0.4 nm to 1.2 nm at a spacing of 0.05 nm for the model Lennard-Jones polymer. Each of the umbrella sampling simulations started with configuration corresponding to desired radius of gyration, as obtained from the aforementioned equilibrium simulations. Then each of the configurations were subjected to a harmonic restraint to ensure a gaussian distribution of the radius of gyration around each desired value of the radius of gyration. Each of the umbrella sampling windows were sampled for 20 ns in NPT ensemble. We used Nose-Hoover thermostat^42,43^ for maintaining the average temperature of 300 K and the Parrinello- Rahman barostat^44^ for maintaining the average pressure of 1 bar. All the water molecules were simulated as rigid molecule using SETTLE algorithm.^45^ Finally, Weighted Histogram Analysis Method (WHAM)^46,47^ was used over the last 10 ns of each of the umbrella sampled trajectories to generate unbiased histograms and the corresponding potentials of mean force or free energies. The total simulation length for all umbrella sampling simulations for each osmolyte solution was 340 ns for the model Lennard-Jones polymer. All simulations were performed using Gromacs 5.0.6 or Gromacs 5.1.4^48^ patched with Plumed^49^ plugin to enable umbrella sampling simulation along the radius of gyration.

All free energy (PMF) calculations have been statistically averaged over at least 4 independent umbrella sampling simulations and free energy difference (Δ*G*^*CE*^= *G*^*E*^ - *G*^*C*^) between collapsed (C) and extended (E) conformations of polymer are compared across different solutions. For a systematic comparison across all osmolytes and to dissect the intrinsic effect of osmolytes from that of neat water, we calculate the change in Δ*G*^*CE*^ of the polymer (hereby referred as ΔΔ*G*) in an aqueous osmolyte solution relative to that in neat water: ΔΔ*G* is defined by

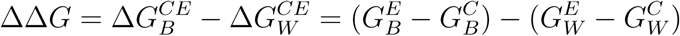

where, B denotes the binary mixtures of water and osmolytes and W denotes the neat water. We employed an experimentally relevant quantity called preferential binding coefficient (Г*s*)_26,27_

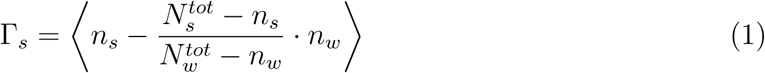

where *n*_*s*_ is the number of cosolutes (glycine or TMAO) bound to the polymer and 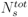 is the total number of cosolutes in the system. On the other hand, *n*_*w*_ is the number of water molecules bound to the polymer and 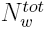 is the total number of water molecules in the system. Г_*s*_ quantifies the excess of cosolute molecules *s* in the polymer solvation shell as compared to its average concentration in the solution. We calculate the profile of Г_*s*_ as a function of distance from the polymer. Towards this end, we employed similar protocol as previously implemented by Mondal et al.^25,28^ To compare the relative preferential binding of each of the cosolutes, we have taken the central atom (O for water, N for TMAO and glycine) of the cosolutes, solvent and any polymer bead for which the distance is shortest. Towards this end, we carry out additional simulations which are as follows: To obtain better statistics, we restrain the polymer either in its respective collapsed or extended conformation of the polymer and then simulated the systems for 15 ns and repeated for thirty independent velocity seeds (hence a total of 450 ns for each of the conformation). Previously,^32^ we have also verified the effect of the magnitude of force used to restrain the polymer in either collapsed or extended conformation by repeating the simulation in the presence of a weak harmonic force along the radius of gyration of polymer. We have found that the qualitative results hold good irrespective of the magnitude of restraints.

## Results and Discussion

The free energetics of the single polymer chain at fixed polymer intramolecular interaction (*∊*_*b*_ = 1.0 kJ/mol) along its radius of gyration in gas phase is known to give rise to a deep free-energy minima at *R*_*g*_ = 0.45 nm corresponding to a collapsed conformation,^25^ which is typical of a hydrophobic polymer. Since the current work aims to dissect the role of the osmolytes on hydrophobic polymer collapse, we mainly explore the effect of variation of polymer-osmolyte dispersion interaction on stabilization of free energetics of polymer-collapse. However, we note that the simultaneous variation of both polymer-water interaction and polymer-osmolyte interaction can potentially mask the intrinsic effect of osmolytes towards the polymer conformation. This is mainly because water itself imparts its denaturing effect by favoring more the extended conformation of polymer over collapsed conformation compared to that in gas phase. As a result, the sole or intrinsic effect of osmolytes on the polymer conformation can not be clearly segregated by simultaneous variation of polymer-osmolyte and polymer-water interaction, as it is not expected that the total effects will be linearly additive.

In this context, for the current investigation, the polymer’s internal bead-bead dispersion interaction (*∊*_*b*_) has been kept fixed at 1.0 kJ/mol (modeled by Lennard Jones interaction) which ensures strong intramolecular interactions and collapse representative of a hydrophobic macromolecules. On the other hand, polymer-water dispersion interaction is kept constant by fixing polymer’s contribution to the Lennard Jones interaction *∊*_*p*_ at 1.0 kJ/mol via com-bination rule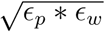 where *∊*_*w*_ is the Lennard Jones interaction of oxygen atoms of SPC/E water model. At this fixed intra-polymer bead-bead interaction and polymer-solvent interaction, we examine the effect of polymer-osmolyte dispersion interaction on its conformational stability by varying the value of *∊*_*p*_ in the polymer-osmolyte combination rule defined by 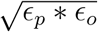, where *∊*_*o*_ represents the interaction of different osmolytes atoms. (However, nevertheless, we will also later briefly show in SI the results obtained by simultaneous variation of polymer-osmolyte and polymer-water interaction.)

Figure 2 A and B depict the free energy profiles of polymer conformations in 0.5 M aqueous glycine and 0.5 M aqueous TMAO solutions respectively, at fixed water-polymer interaction but at three different polymer-osmolyte interactions quantified by *∊*_*p*_ = 0.6, 1.0 and1.4 kJ/mol. In all cases, the free energy profiles of polymer in aqueous osmolyte solutions are also compared with that in neat water (i.e. in absence of cosolutes). The overall shape of the free energy profile of this polymer has been elaborately discussed in previous works by Mondal et al.^25,32^ Figure 2 A and B suggests that the polymer’s conformational equilibrium is majorly dictated by Δ*G*^*CE*^ (= *G*^*E*^ - *G*^*C*^) i.e. the relative free energetic (de)stabilization of collapsed conformation at *R*_*g*_ ≈ 0.45 nm relative to the extended or unfolded polymer conformation at *R*_*g*_ ≈ 1.10 nm. We find that, at *∊*_*p*_ = 0.6 kJ/mol, the extent of stabilization of the collapsed conformation of the polymer compared to extended conformation is higher in both aqueous solution of glycine and aqueous solution of TMAO compared to that in neat water (having same polymer-water interaction). Thereby, at *∊*_*p*_ = 0.6 kJ/mol, when polymer-water interaction is fixed in all solution, both osmolytes act as a protectant of collapsed conformation with similar degree of protection. However, with systematic increase of polymer-glycine interaction strength from *∊*_*p*_ = 0.6 kJ/mol to 1.4 kJ/mol at an interval of kJ/mol, the extent of stabilization of collapsed conformation over extended conformation monotonically decreases relative to that in neat water. In other words, with gradual increase of polymer-osmolyte interaction, aqueous solution of glycine changes its role from being a protectant to a denaturant of collapsed conformation. On the contrary, TMAO, with increasing degree of polymer-osmolyte interactions but at fixed polymer-water interaction, monotonically increases the extent of stabilization of collapsed conformation relative to extended conformation and hence manifests an increasing degree of protection.

**Figure 2:**
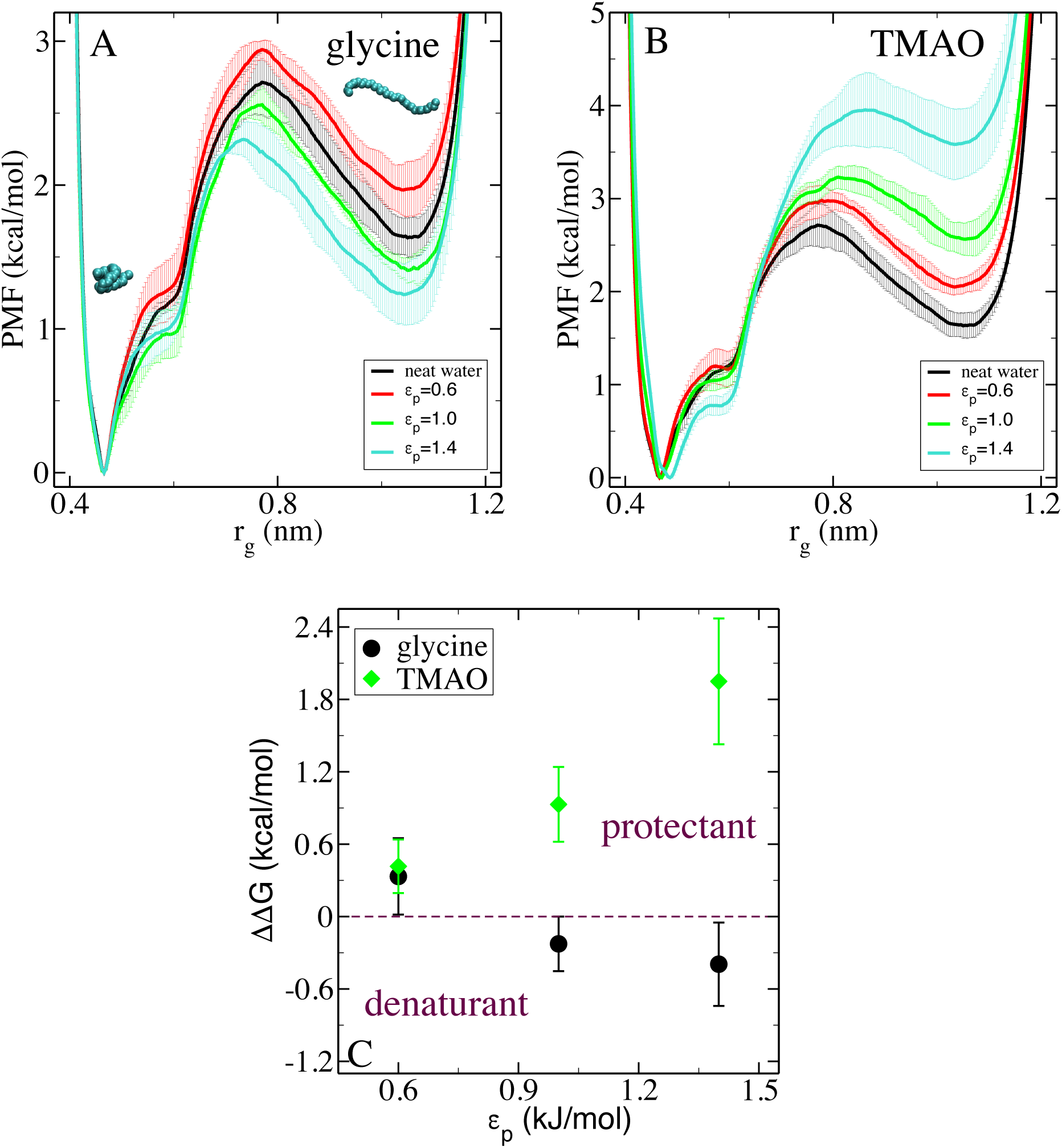
Potential of mean force (PMF) or free energy of the polymer along the radius of gyration in pure water (A or B) and in two aqueous osmolytes solutions (glycine (A) and TMAO (B)) for different polymer osmolytes interactions (*∊*_*p*_) keeping polymer-water interaction (*∊*_*p*_ = 1.0 kJ/mol) fixed.(C) Change in Δ*G*^*CE*^ of the polymer in aqueous solutions of glycine and TMAO with respect to neat water (ΔΔ*G*) along different polymer-cosolutes interactions (*∊*_*p*_) keeping polymer-water interaction (*∊*_*p*_ = 1.0 kJ/mol) fixed.

The progression of ΔΔ*G* (given by 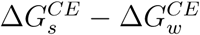 where *s* refers to aqueous osmolyte solution and *w* refers to neat water) as a function of polymer-osmolyte interaction (*∊*_*p*_) (see Figure 2 C) provides a quantitative measure of these observations at a fixed polymer-water interaction. A benefit of using ΔΔ*G* in the analysis is that the polymer’s own intrinsic effect towards free energetics gets cancelled out upon subtraction and we get a measure of osmolyte’s intrinsic effect towards stabilization of polymer conformation. We find that ΔΔ*G* value is always positive in case of TMAO and monotonically increases with increase in *∊*_*p*_, confirming the protecting role of TMAO across variation of polymer-osmolyte interactions. On the other hand, ΔΔ*G* value is positive for glycine at a smaller value of *∊*_*p*_ = 0.6 kJ/mol, suggesting its role as a protectant at low *∊*_*p*_. However, with further increase in *∊*_*p*_ to 1.4 kJ/mol through 1.0 kJ/mol, ΔΔ*G* value becomes gradually negative for glycine illustrating its crossover from being a protectant at lower polymer-osmolyte interaction to a denaturant at higher polymer-osmolyte interaction. As an aside, we here note that we have also investigated the role of osmolytes by simultaneously varying both polymer-water and polymer-osmolyte interaction using geometric combination rule. As illustrated in Figure S1 in SI, the overall trends as discussed earlier also remain qualitatively similar. However, for the purpose of capturing the intrinsic effects of osmolytes, all future discussions hereby will involve the investigations using fixed polymer-water interaction.

The three-dimensional density iso-surfaces of these two cosolutes around the polymer illustrate (Figure 3) a qualitative picture of the cosolute’s interaction with the polymer surface at varying *∊*_*p*_. At very high *∊*_*p*_ of 1.4 kJ/mol studied in this current work, both osmolytes can favorably bind to the polymer surface. However, glycine tends to exclude from the surface of both collapsed and extended conformation at smaller *∊*_*p*_ of 0.6 and 1.0 kJ/mol. On the other hand, TMAO preferably accumulates near polymer surface even at smaller values of *∊*_*p*_ of 0.6 and 1.0 kJ/mol, although the extent of TMAO population increases with *∊*_*p*_. As we would see later, the contrasting picture of TMAO’s interaction with the polymer compared to that of glycine would play a decisive role in regards to polymer’s contrasting collapse behavior in presence of these two osmolytes. As will be discussed in SI text and demonstrated in Figure S2, the overall behavior is consistent with polymer-solute or polymer-solvent radial distribution function.

**Figure 3:**
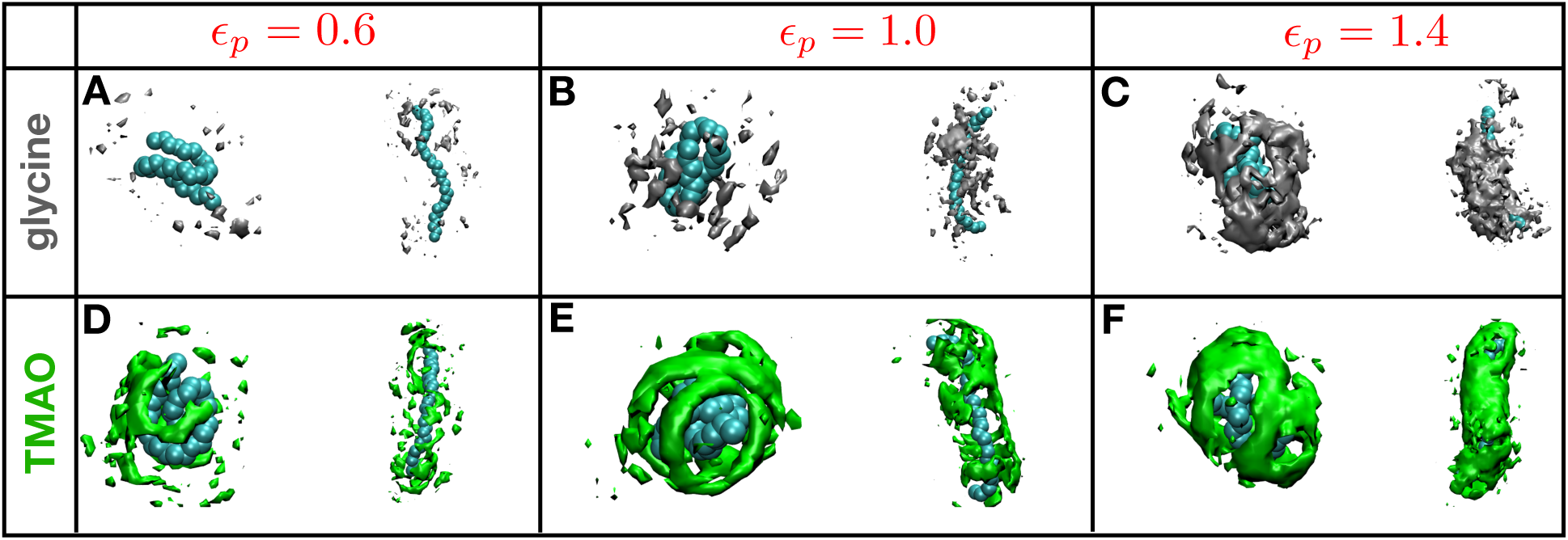
Density isosurfaces within 1 nm from the collapsed and extended polymer chain for different polymer-osmolyte interactions (*∊*_*p*_, in kJ/mol), keeping polymer-water interaction (*∊*_*p*_ = 1.0 kJ/mol) fixed, in aqueous solutions of glycine (A,B,C) and TMAO (D,E,F) (isovalue is identical for all cases).

A systematic approach towards understanding the aforementioned behavior of the hydrophobic polymer in different osmolyte solutions is the Wyman-Tanford theory of preferential solvation. The Wyman-Tanford approach hypothesizes the preferential binding coeffi-cient r_*s*_ (see equation 1)^26,27^ as the key quantity to describe the dependence of the polymer conformational equilibria on cosolutes. As discussed in method, r_*s*_ quantifies the excess of cosolute molecules *s* in the polymer solvation shell as compared to its average concentration in the solution.

Г_*s*_ *¾* 0 if the osmolyte preferentially accumulates next to the polymer and is < 0 if it is excluded from the polymer. Note that r_*s*_ is closely related to a frequently used parameter known as Kirkwood Buff integral.^24,50^

The effect of preferential binding on a conformational equilibrium between the collapsed and the extended configurations C ⇌ E (with an equilibrium constant *K*) is usually interpreted in terms of the thermodynamic analysis put forward by Wyman and Tanford,^26,27^ which leads to

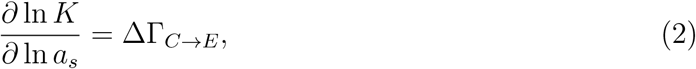

where *a*_*s*_ is the activity of the cosolute in the binary solution. According to Eq. 2, an increase in the concentration of the cosolute would lead to the macromolecular unfolding if Δr > 0, and in contrast would favor the folded state over the unfolded one if Δr <0.

To get molecular insight of binding of different osmolytes and water molecules with the polymer, we calculate preferential binding coefficient (Г_*s*_) of different osmolytes with the collapsed and extended state of the polymer. Figure 4A,B illustrates the profile of preferential binding of both osmolytes on the collapsed and extended conformation of the hydrophobic polymers as a function of distance from the polymer surface (see SI text for detailed method). As observed in the case of glycine (Figure 4A), at an weaker polymer-osmolyte interaction (*∊*_*p*_ = 0.6 kJ/mol), Г_*s*_ of glycine towards both collapsed and extended conformation of the polymer is negative, indicating that at a weaker polymer-osmolyte interaction, glycine molecules will be preferentially excluded from the polymer surface, compared to water molecules. However, as shown in Figure 4D, we find that, glycine molecules are predicted to exclude more from an extended polymer surface than from the collapsed polymer surface so that Δr < 0 for *∊*_*p*_ = 0.6 kJ/mol. Thereby, as put forward by Wyman-Tanford approach,glycine will act as an osmo-protectant for a hydrophobic polymer at an weaker *∊*_*p*_. This prediction is in perfect agreement with relevant polymer free energy profile depicted earlier in Figure 2A and Figure 2C for glycine. The observation of preferential exclusion of glycine from polymer surface at weaker polymer-glycine interaction is reminiscent of what has been previously perceived on glycine’s protectant role of protein’s native conformation.^19^ On the contrary, at increasing polymer-glycine interactions (and keeping polymer-water interaction fixed as before), Г_*s*_ > 0 for glycine for both collapsed and extended conformation of the polymer, implying that glycine molecules start getting preferentially bound to the polymer surface at higher *∊*_*p*_. However, intriguingly, from Figure 4C, we find that Δr *>* 0 for *∊*_*p*_ = 1.0 and 1.4 kJ/mol i.e. more glycine molecules would preferentially bind to extended conformation of the polymer compared to collapsed conformation, thereby implying, as per equation 2, a cross-over of glycine’s role as an osmoprotectant at weaker polymer interaction to a denaturant at stronger polymer interaction. This preferential binding based argument of glycine’s action towards polymer surface,especially the cross-over, is in perfect agreement with the conclusions drawn on the basis of free energy landscape of polymer (see Figure 2) in aqueous solution of glycine.

**Figure 4:**
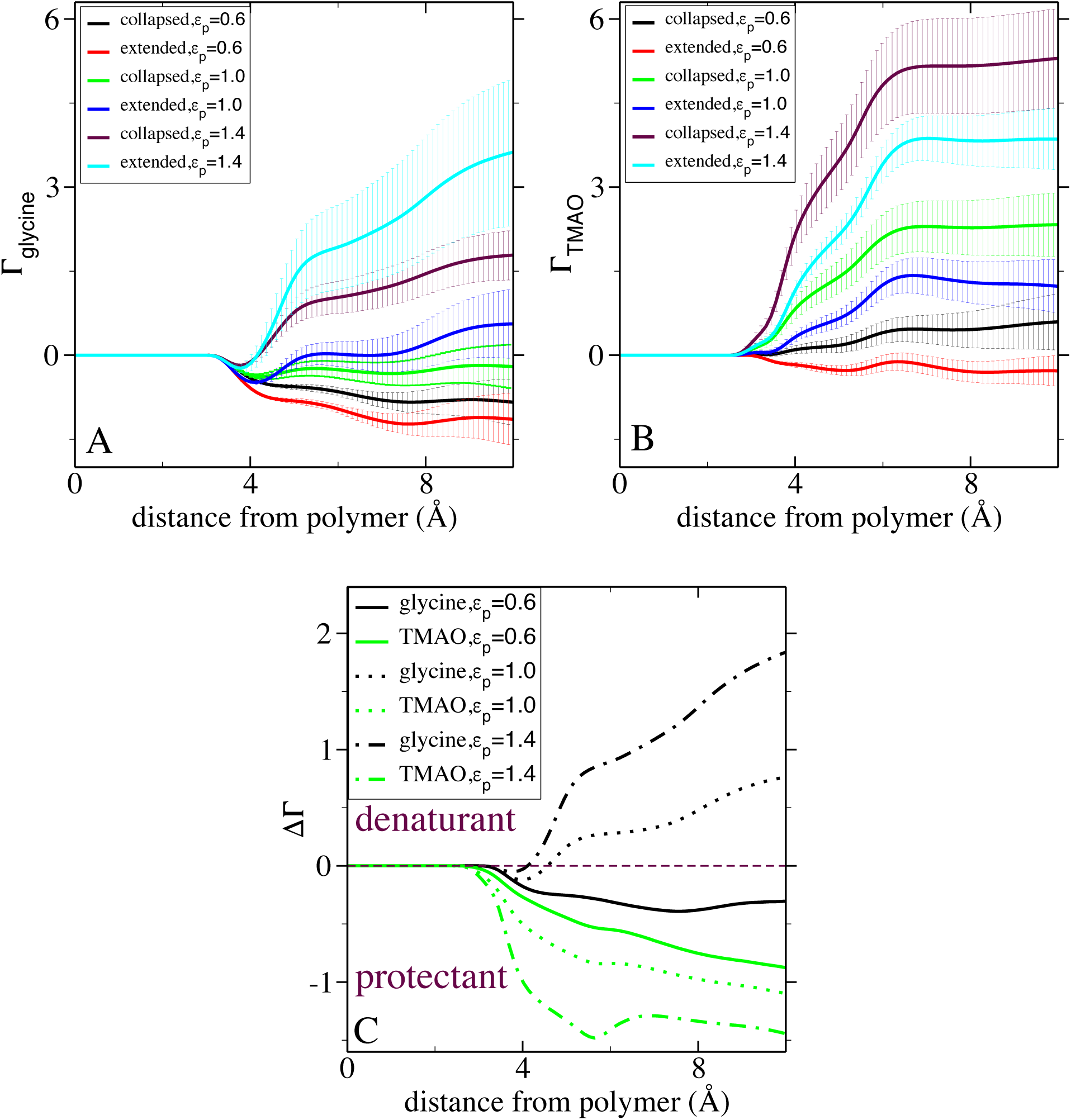
The preferential binding coefficient (r) along the distance from the polymer at varying polymer-osmolyte interactions (*∊*_*p*_), keeping polymer-water interaction (*∊*_*p*_ = 1.0 kJ/mol) fixed, in aqueous solution of glycine (A) and TMAO (B) and the differences Δr(errorbars are not shown for visualization purposes) (C) along the distance from the polymer.

Aqueous solution of TMAO displays quite distinct picture of conformation-dependent preferential interaction compared to that of glycine. As observed in Figure 4B, TMAO molecules preferentially bind to polymer surface over water molecule almost across all *∊*_*p*_, implying Г_*s*_ > 0. However, across all *∊*_*p*_, extent of preferential binding of TMAO molecules is relatively higher on the collapsed conformation of polymer surface than on its extended conformation making Δr < 0 as shown in Figure 4C. This proposition based on the molecular theory of preferential interaction of TMAO is in unison with free energy profile of this polymer in presence of TMAO illustrated earlier in Figure 2B and Figure 2C. The overall results robustly unify TMAO’s effect for polymer studied independently by Mondal et al^25^ and for protein investigated by Cremer and coworkers.^19^ Cremer and coworkers^19^ have experimentally and computationally showed that while acting as a protecting agent, TMAO can still bind to the protein surface. Similar observation has also been previously made by Mondal et al^25^ regarding TMAO’s effect on a hydrophobic polymer where it was found that it is the relatively higher preferential binding of TMAO molecules to the collapsed conformation of polymer than that of extended conformation that swings the hydrophobic polymer’s conformational equilibrium in favor of collapsed state. Overall results clearly demonstrate the origin of mutually contrasting behaviors of two protein-stabilizing osmoprotectants towards hydrophobic interaction.

## Conclusion

Figure 5 concisely summarizes the overall picture of both osmolytes’ role on controlling hydrophobic polymer conformation at a wide range of polymer-cosolute interaction (for a given polymer-water interaction), in consonance with Wyman-Tanford theory of preferential interaction. The illustration of the relative trend of ΔΔ*G* and Δr for glycine unequivocally suggests a clear cross-over of glycine’s role from being an osmoprotectant at smaller polymer-cosolute interaction via principle of preferential exclusion to a denaturant at larger polymer-cosolute interaction via principle of preferential binding. Fig. 5 also represents TMAO’s consistent trend as an osmoprotectant of polymer’s collapsed conformation via principle of preferential binding at wider range of polymer-cosolute interaction.

**Figure 5:**
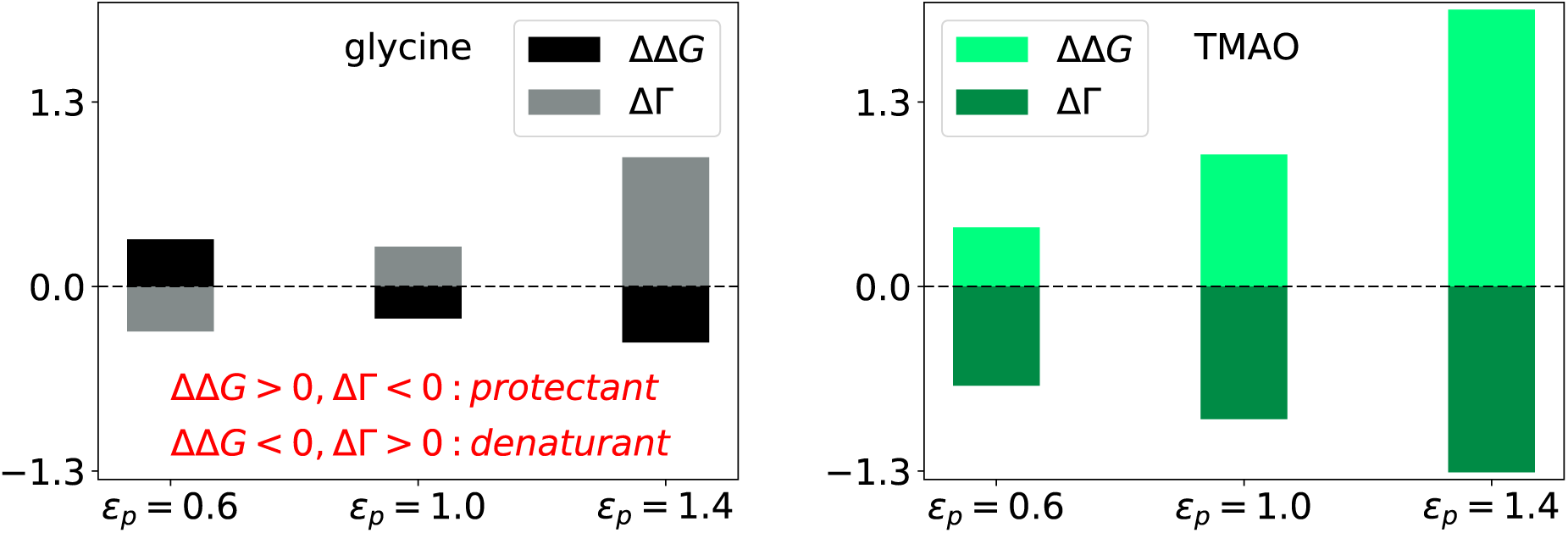
ΔΔ*G* and Δr in aqueous solution of glycine and TMAO at different polymerosmolyte interactions (*∊*_*p*_, in kJ/mol) keeping polymer-water interaction (*∊*_*p*_ = 1.0 kJ/mol) fixed (for calculating Δr we have used first solvation shells of polymer in different osmolytes.)

For comparison purpose, in SI text, we also provide a similar analysis using a third osmolyte namely glycine betaine, subject of many previous works. Figure S3 illustrates the overall effect of glycine betaine and its preferential interaction for weaker and stronger polymer-osmolyte interactions. We find that the effect of glycine betaine on polymer collapse is very similar to that of glycine: a cross-over from being a protectant at weaker polymer-osmolyte interaction to a being denaturant at stronger polymer-osmolyte interaction is mainly triggered by the reverse trend in relative preferential interaction of glycine betaine towards collapsed and extended conformation.

At this stage, a qualitative chemical insight can only be offered in support of the these observations. We believe that TMAO’s consistent preferential binding with the collapsed polymer conformation relative to extended conformation is attributed to TMAO’s amphiphilic nature. The hydrophobic methyl group in TMAO favorably binds with the polymer’s collapsed conformation than the extended conformation due to availability of local hydrophobic contacts in the former. This preferential binding of TMAO with collapsed conformation is reinforced with increasing polymer-osmolyte interaction which promotes the protectant behavior of TMAO. On the other hand, glycine lacks any hydrophobic functional groups unlike TMAO. Hence, at weak polymer-osmolyte interaction, the polar nature of glycine excludes any contact with polymer, especially with extended conformation and hence the collapsed conformation of polymer is promoted at weak polymer-osmolyte interaction in aqueous glycine solution. However, with increasing polymer-osmolyte interaction, glycine eventually binds to polymer but this polymer-glycine binding is not driven by hydrophobic interaction. Rather the binding is promoted more with extended conformation due to larger access of surface area. However, future work is certainly needed for further validation of this hypothesis.

The formation of a hydrophobic core is crucial to the protein folding which leads to function of most proteins in the cell. The distinct and contrasting trends of the osmolytes displayed in the current work in dictating the conformation of hydrophobic polymer are found to be statistically significant and can have implication in stabilization of hydrophobic interactions. The observations of these osmolytes’ mutually contrasting role towards hydrophobic interaction are quite distinct from both of the osmolytes’ uniform stabilising role towards protein. The results discussed in the current work unify different spectrum of previous works of osmolyte’s effects of protein by Cremer and co-workers^19^ and that on hydrophobic system by Mondal et al^25,32^ on similar ground. Considering the fact that the proteins’ topology is usually constituted by both hydrophobic and hydrophilic groups, our current results call for a quantitative dissection of osmolytes’ effect on proteins’ different segments. We believe that an investigation of effect of these osmolytes towards macromolecule’s electrostatic interaction and topology certainly holds a promising future prospect.

## Acknowledgement

This work was supported by computing resources obtained from shared facility of TIFR Center for Interdisciplinary Sciences, India. JM would like to acknowledge research intramural research grants obtained from TIFR, India, Ramanujan Fellowship and Early Career Research funds provided by the Department of Science and Technology (DST) of India (ECR/2016/000672). Part of the work was carried out in San Diego supercomputing resources provided by XSEDE (TG-CHE160057) to JM.

## Supporting Information (SI)

Graphs illustrating free energetics by varying both polymer-osmolyte and polymer-water interaction, polymer-osmolyte radial distribution function, free energetics in case of glycine betaine (PDF).

## Supporting information for “Heterogeneous Impacts of Protein-Stabilizing Osmolytes on Hydrophobic Interaction”

### Quantification of polymer-osmolyte pair correlation

Fig. S2 in SI text provides a quantitative account of the polymer’s interaction with the cosolutes and water by radial distribution function (RDF). At smaller values of *∊*_*p*_ of 0.6 and 1.0 kJ/mol, the RDFs between polymer and glycine are significantly smaller than unity, implying that glycine remains relatively depleted from the polymer surface at weaker polymer-glycine interaction. On the other hand, even at comparable values of *∊*_*p*_, as quantified by Fig. S2 in SI text, TMAO interacts favorably with the polymer surface. We also find that at very high value of *∊*_*p*_ of 1.4 kJ/mol, the interaction of all cosolutes with the polymer becomes favorable. Interestingly, these polymer-cosolute pair correlation profiles also display considerable dependence on polymer-conformations (collapsed or extended). As illustrated in Fig. S2, TMAO displays higher correlation with collapsed conformation of the polymer compared to that with extended conformation across entire range of *∊*_*p*_ investigated in this work. Quite intriguingly, for the similar polymer-osmolyte interaction range, glycine shows a reverse trend on its respective correlation with polymer conformation: the value of pair-correlation being higher with extended conformation than that with collapsed conformation of polymer. As we will discover in the following section, these relative conformation-dependences of polymer-cosolute preferential interaction would play decisive roles in guiding the polymer conformation equilibrium.

---

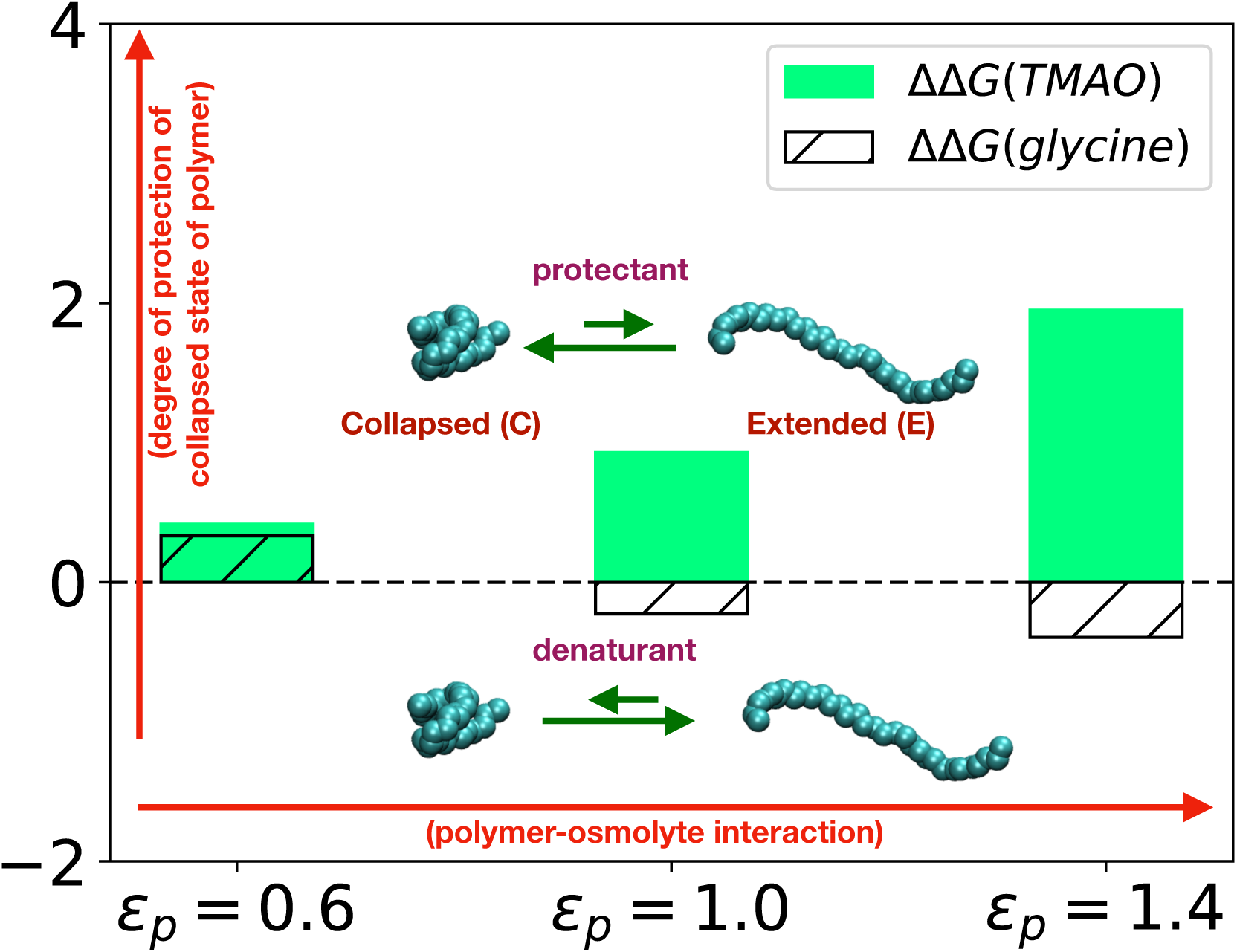

**Figure S1:**
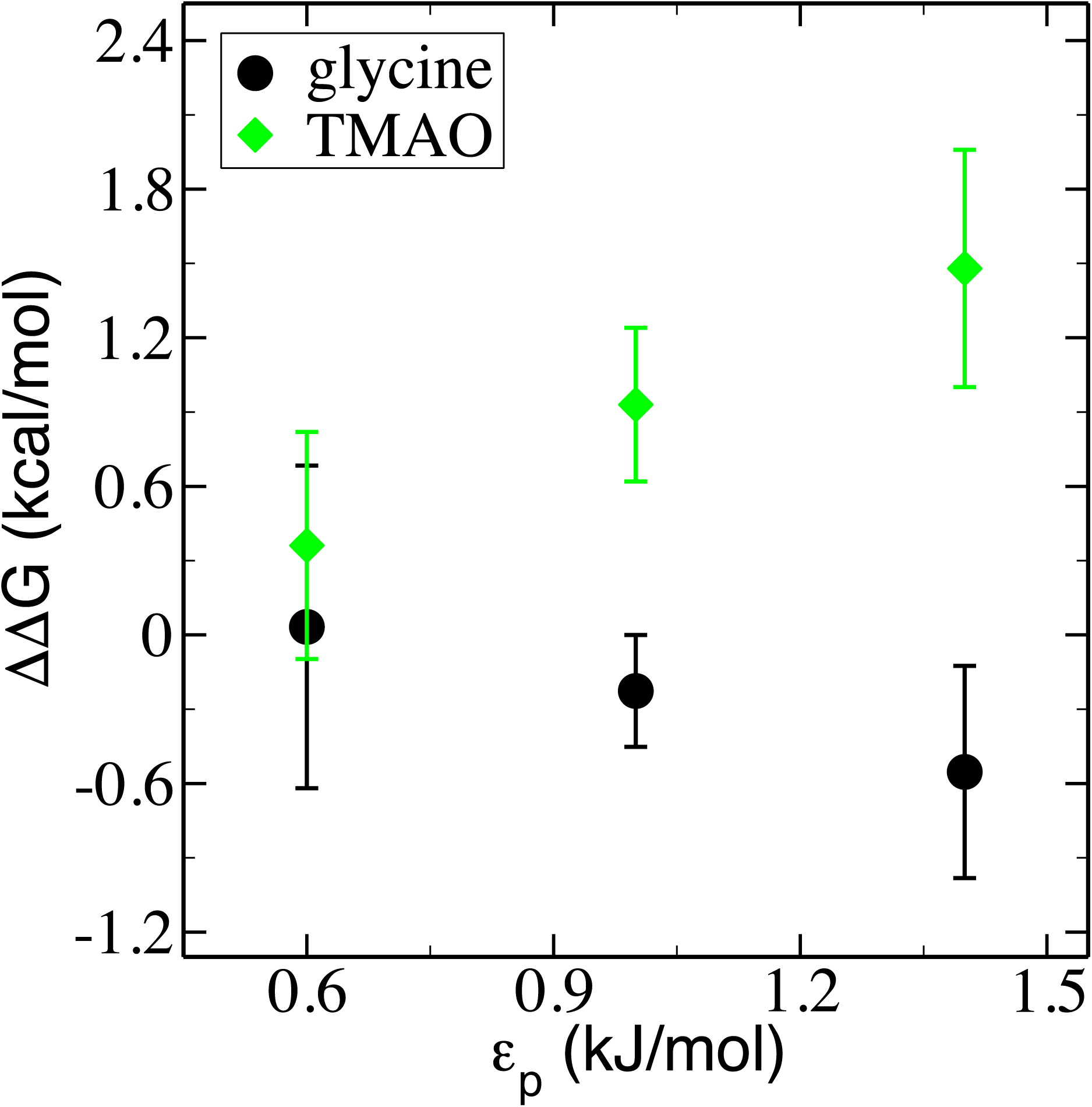
Change in L’*G*^*CE*^ of the polymer in aqueous solutions of glycine and TMAO with respect to neat water (L’L’*G*) along different polymer-cosolutes interactions (*∊*_*p*_) and different polymer-water interaction.

**Figure S2:**
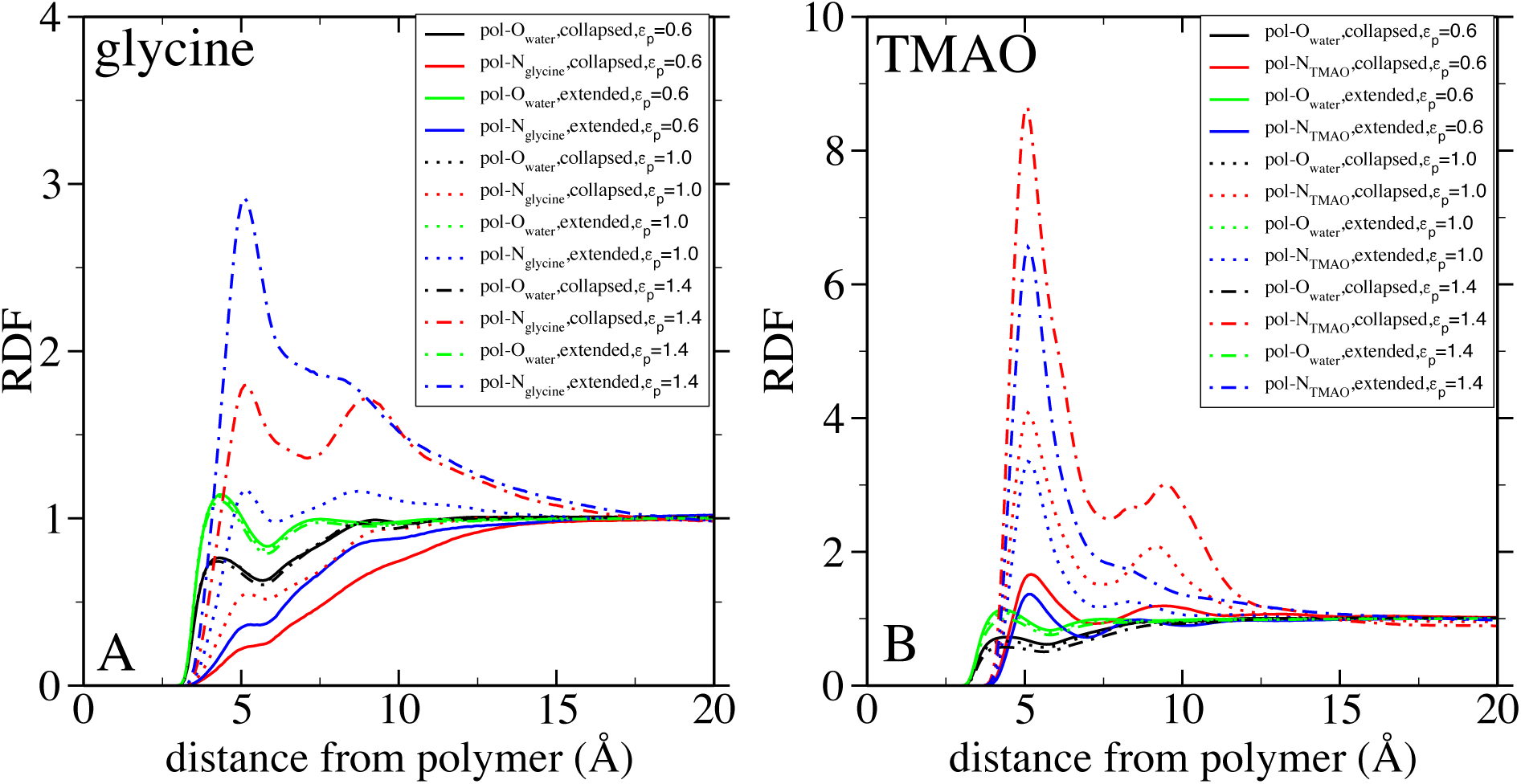
The radial distribution functions (RDF) between any polymer (collapsed or extended) bead and O of water (A or B) and polymer bead-central atom of glycine (A) and TMAO (B) for different polymer-osmolyte interactions (*∊*_*p*_) keeping polymer-water interac-tion (*∊*_*p*_ = 1.0) fixed. Here we have chosen the central atom as N for the osmolytes.

**Figure S3:**
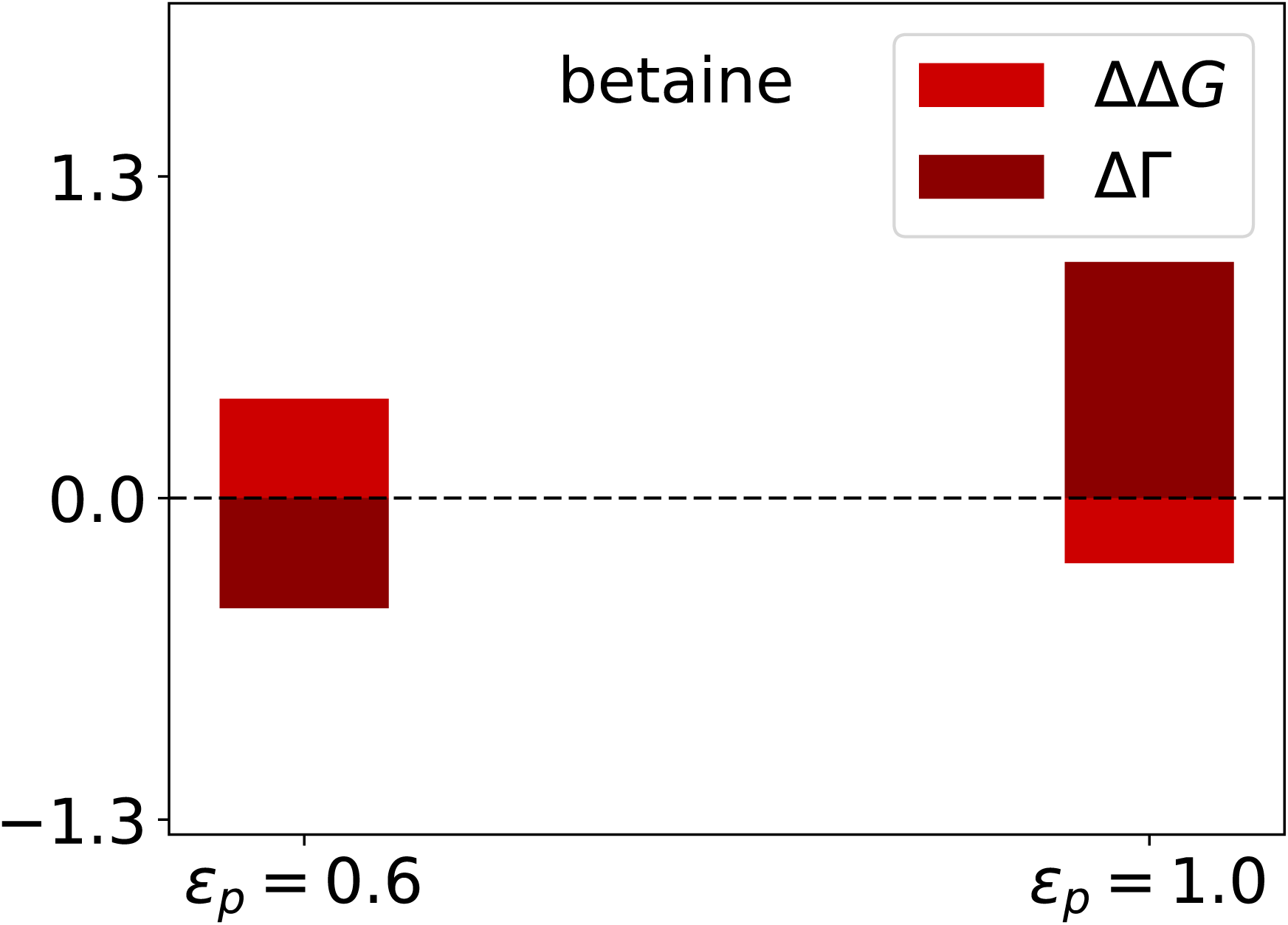
L’L’*G* and L’r in aqueous solution of glycine betaine at different polymer-osmolyte interactions (*∊*_*p*_) keeping polymer-water interaction (*∊*_*p*_ = 1.0) fixed (for calculating L’r we have used first solvation shells of polymer in different osmolytes.) Force field parameters for glycine betaine were adopted from Cremer and coworkers^1^

